# Maternal loading of a small heat shock protein increases embryo thermal tolerance in *Drosophila melanogaster*

**DOI:** 10.1101/150102

**Authors:** Brent L. Lockwood, Cole R. Julick, Kristi L. Montooth

## Abstract

Maternal investment is likely to have direct effects on offspring survival. In oviparous animals whose embryos are exposed to the external environment, maternal provisioning of molecular factors like mRNAs and proteins may help embryos cope with sudden changes in the environment. Here we sought to modify the maternal mRNA contribution to offspring embryos and test for maternal effects on acute thermal tolerance in early embryos of *Drosophila melanogaster*. We drove in vivo overexpression of a small heat shock protein gene (*Hsp23*) in female ovaries and measured the effects of acute thermal stress on offspring embryonic survival and larval development. We report that overexpression of the *Hsp23* gene in female ovaries produced offspring embryos with increased thermal tolerance. We also found that brief heat stress in the early embryonic stage (0 to 1 hour-old) caused decreased larval performance later in life (5 to 10 days-old), as indexed by pupation height, as well as increased development time to pupation. Maternal overexpression of *Hsp23* protected embryos against these heat-induced larval defects. Our data demonstrate that transient products of single genes have large and lasting effects on whole-organism environmental tolerance. Further, our results suggest that maternal effects have a profound impact on offspring survival in the context of thermal variability.

**SUMMARY STATEMENT:** A gene-specific maternal effect confers thermal tolerance to offspring embryos in the fruit fly *Drosophila melanogaster*.

## INTRODUCTION

Acute thermal stress is principally felt at the cellular and biochemical levels through the disruption of macromolecular structures (Richter et al., 2010; Somero et al., 2017). These thermal perturbations pose challenges for ectotherms that live in variable thermal environments where sudden changes in temperature are a frequent occurrence (Denny et al., 2011; Terblanche et al., 2011; Dowd et al., 2015; Buckley and Huey, 2016). Thermal stress causes proteins to unfold, which not only leads to the loss of protein function but also to protein aggregation that is toxic to cells (Richter et al., 2010; Somero et al., 2017). To combat these effects, nearly all living organisms possess a conserved set of cellular responses—collectively referred to as the heat shock response or cellular stress response—which are characterized by rapid shifts in the expression of hundreds to thousands of gene loci (Gasch et al., 2000; Leemans et al., 2000; Buckley et al., 2006; Lockwood et al., 2010; Brown et al., 2014). A key component of the heat shock response is the dramatic induction of genes that encode heat shock proteins (HSPs), while the majority of the rest of the proteome ceases to be expressed (Tissiéres et al., 1974; Mirault et al., 1978; Lindquist, 1981; Hofmann and Somero, 1996; Tomanek and Somero, 1999; Tomanek and Zuzow, 2010). HSPs function as molecular chaperones that bind, sequester, and help refold thermally denatured proteins (Richter et al., 2010), providing thermal protection at the molecular level that scales up the whole organism. Indeed, sublethal thermal exposures that induce the heat shock response allow organisms to survive more extreme subsequent thermal exposures that would otherwise be lethal (Arrigo, 1987; Feder et al., 1996). Transgenic overexpression of HSPs confers increased whole-organism thermal tolerance (Welte et al., 1993; Feder et al., 1996), and the expression of HSPs has been shown to be adaptive under conditions of heat stress, as laboratory selection to high temperatures leads to higher expression of HSPs (Rudolph et al., 2010). In addition, many populations and species that live in environments characterized by frequent, acute exposures to extreme heat, have evolved higher expression of HSPs than closely related species that inhabit more benign thermal environments (Hofmann and Somero, 1996; Tomanek and Somero, 2000; Dong et al., 2008; Lockwood et al., 2010; Schoville et al., 2012; Dilly et al., 2012).

Despite the broad evolutionary conservation of the heat shock response across taxa (Kültz, 2005; Somero et al., 2017), animals in the earliest life stages have vastly reduced heat shock responses (Graziosi et al., 1980; Welte et al., 1993) due to the low transcriptional activity of early zygotes (Tadros and Lipshitz, 2009). This poses a challenge to oviparous species with external embryonic development. Early embryos of these organisms are directly exposed to the thermal environment and may have little opportunity to express protective proteins from their own genomes. Rather, their mechanisms for coping with thermally-induced molecular damage are limited to the molecular factors (i.e., RNAs, protein, and organic osmolytes) that are loaded into eggs by mothers (Wieschaus, 1996). Indeed, previous studies have shown early embryonic stages to be more thermally sensitive than later stages (Walter et al., 1990; Welte et al., 1993).

Given that maternal oogenesis establishes the early embryonic transcriptome and proteome (Schüpbach and Wieschaus, 1986; Wieschaus, 1996; Tadros and Lipshitz, 2009), maternal molecular factors are likely to be a major determinant of developmental robustness and survival in the face of variable thermal environments. However, few studies have characterized the molecular roles of maternal effects in the context of embryonic thermal tolerance (Sato et al., 2015). In fruit flies (*Drosophila melanogaster*), the early time window of minimal zygotic transcriptional activity spans the first 2 hours of development, after which zygotic transcription begins to predominate over the maternally provided pool of mRNAs (Blythe and Wieschaus, 2015a). Consequently, 4-h-old embryos are more heat tolerant than earlier stages (Walter et al., 1990), and by 14 h post-fertilization (Welte et al., 1993) embryos attain approximately the same degree of thermal tolerance that they possess later on as larvae, pupae, and adults (Huey et al., 1991; Feder et al., 1997).

Among the mRNAs that are loaded into eggs by *D. melanogaster* mothers, messages that encode members of the small heat shock protein (sHSP) family are highly abundant, with two sHsps being among the top 1% of most highly abundant transcripts in the early embryo (see Fig. 1)(Pauli et al., 1989; Michaud and Tanguay, 2003; Brown et al., 2014; Morrow and Tanguay, 2015). Genes encoding this class of proteins are also among the most highly expressed genes following heat stress in larvae, pupae, and adults (Berger and Woodward, 1983; Ayme and Tissières, 1985; Horwitz, 1992; Brown et al., 2014). sHSPs are a family of molecular chaperones that serve a wide-range of molecular functions, including stabilizing major cellular structural components like the cytoskeleton (Leicht et al., 1986; Horwitz et al., 1992) and the cell membrane (Tsvetkova et al., 2002; Horváth et al., 2008). Thus, the maternal contribution of these proteins may be a critical factor in maintaining embryonic development of offspring in both benign thermal conditions and in the presence of thermal stress.

**Figure 1.**
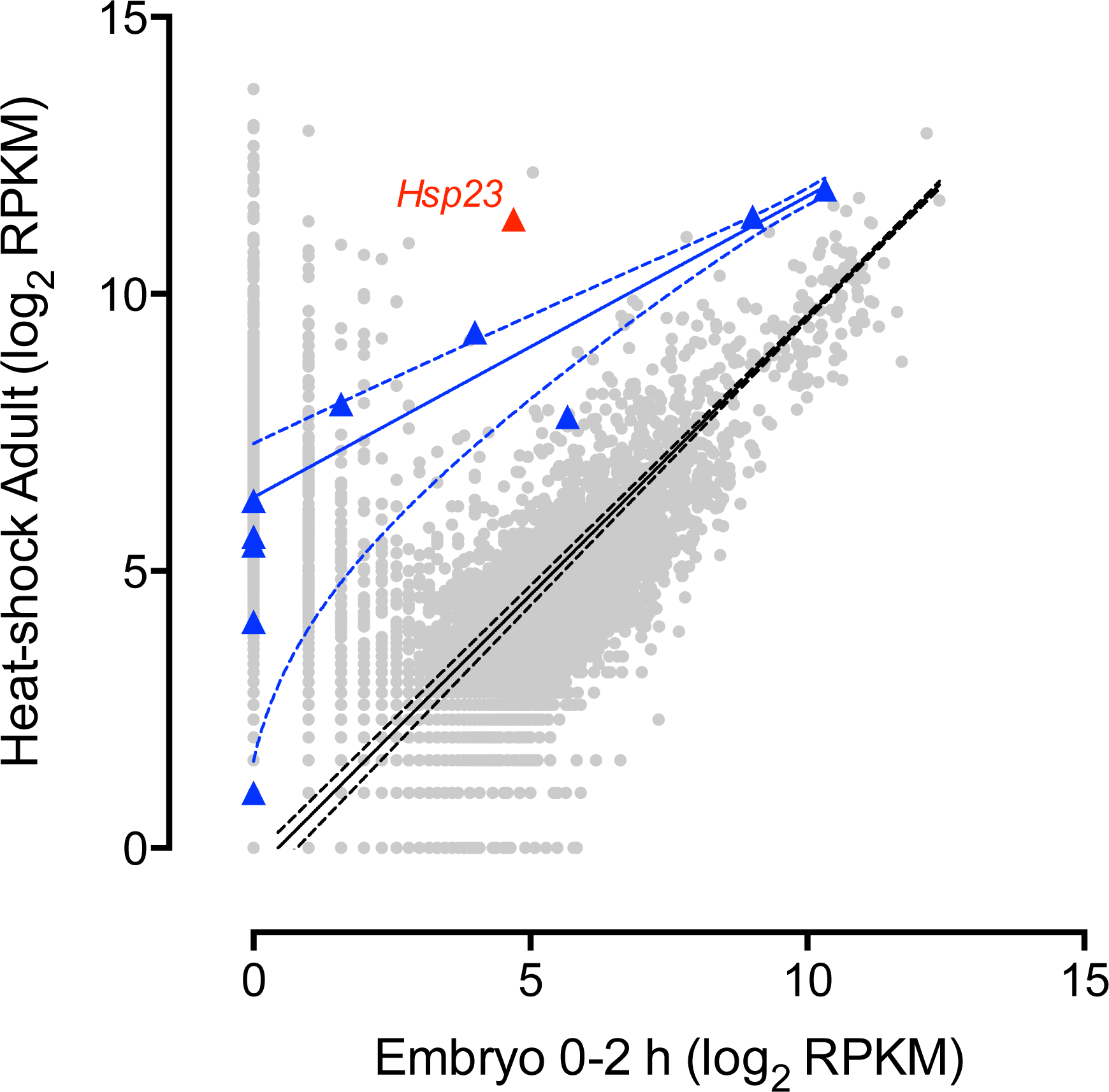
Relationship between transcriptomes and sHSP mRNA levels in early embryos vs. heat-shocked adults. Data represent 18,029 unique transcripts and are expressed as mean expression values (RPKM) on a log_2_-scale of 0-2 hour-old embryos and 4 day-old adults (males and females pooled). Transcripts encoding small heat shock proteins (sHSPs) are shown in blue triangles, *Hsp23* highlighted in red, and all other transcripts are shown in grey circles. The solid black line (+/- 95% confidence bands in black dashed lines) represents the least-squares regression fit of all transcripts (*R^2^* = 0.17, *y* = 10^(1.002*log(*x*) - 0.1344)) and the solid blue line (+/- 95% confidence bands in blue dashed lines) represents the robust regression fit of sHSP transcripts (*R^2^* = 0.97, *y* = 10^(0.5436*log(*x*) + 1.905)), for which *Hsp23* was a significant outlier (ROUT outlier analysis, Q = 1%).

Here, we establish a role for maternal effects in conferring embryonic thermal tolerance in *D. melanogaster* via maternal loading of the sHSP gene *Hsp23*, which is a major component of the heat shock response. We report that among sHSP genes, *Hsp23* is unique in that it is a major component of the adult heat shock response but only present at low abundance in early embryos. Further, by driving overexpression of this gene in female oocytes, we observed marked increases in thermal tolerance in offspring embryos and lasting effects that influenced larval performance—both of which were significant maternal effects. Overall, our results demonstrate that single genes of large effect can contribute significantly to whole-organism phenotypes, such as thermal tolerance, and that maternal loading of mRNAs can influence not only early embryonic development but also larval performance later in life.

## MATERIALS AND METHODS

### modENCODE expression data

modENCODE is a collaborative project that generated transcriptomic data from RNA-sequencing (RNA-Seq) across life stages and in response environmental stressors in *D. melanogaster* (Brown et al., 2014). Expression data were downloaded from FlyBase (Attrill et al., 2016) and consist of mRNA levels (expressed as reads per kilobase of transcript per million mapped reads: RPKM) of 18,029 unique transcripts. Among these transcripts, we used nonlinear least-squares regression fitting to compare mRNA levels in early embryos (0-2 h post-fertilization) and 4 d-old heat-shocked adults (36°C for 1 h), with Robust regression and Outlier removal (ROUT) analysis (Motulsky and Brown, 2006) to identify outliers.

### Fly stocks

To assess the effects of targeted overexpression and increased maternal loading of *Hsp23* in early embryos, we used the Gal4-UAS system (Brand and Perrimon, 1993; Duffy, 2002) in a two-step crossing scheme (Fig. 2A). First, we used a female germline Gal4 driver, *MTD-Gal4* (Bloomington Stock—BL#: 31777), crossed with *UAS-Hsp23* (BL#: 30541) to cause *Hsp23* overexpression in female ovaries (*Hsp23^OE^*). These constructs when brought together in a genetic cross drive overexpression of the target gene (*Hsp23*) in female ovaries and were chosen to modify the levels of *Hsp23* mRNAs that are loaded into eggs. Second, we tested the effects of this overexpression construct in early embryos by comparing the phenotypes of 0-1 h-old offspring embryos from reciprocal crosses between the *Hsp23^OE^* and the control genotype (*w^1118^*) that switched the female and male genotypes, such that embryos from one cross (female *Hsp23^OE^* x male *w^1118^*^)^ had mothers that overexpressed *Hsp23* and embryos from the other cross (female *w^1118^* x male *Hsp23^OE^*) were genetically similar but had control mothers with wildtype *Hsp23* expression. The control genetic background was *w^1118^* (BL#: 5905), which was the original strain used to generate the *UAS-Hsp23* transgenic line. We note that the *MTD-Gal4* strain was generated in the *w\** genetic background. Therefore, F2 offspring of *Hsp23^OE^* mothers received mitochondria from *w\**, whereas offspring from the reciprocal cross received mitochondria from *w^1118^*. While these represent two distinct mtDNA genetic backgrounds, many lab stocks were originally derived from similar mitochondrial lineages and natural populations of *D. melanogaster* harbor relatively low levels of mtDNA polymorphism (Cooper et al., 2015). Thus, we interpret measurable differences in embryonic thermal tolerance among genotypes to be largely the result of differential maternal loading of *Hsp23* mRNAs, and not an artifact of mitochondrial lineage. All stocks were obtained from the Bloomington Drosophila Stock Center (Bloomington, IN) and maintained at 22°C on standard cornmeal, yeast, and agar medium.

**Figure 2.**
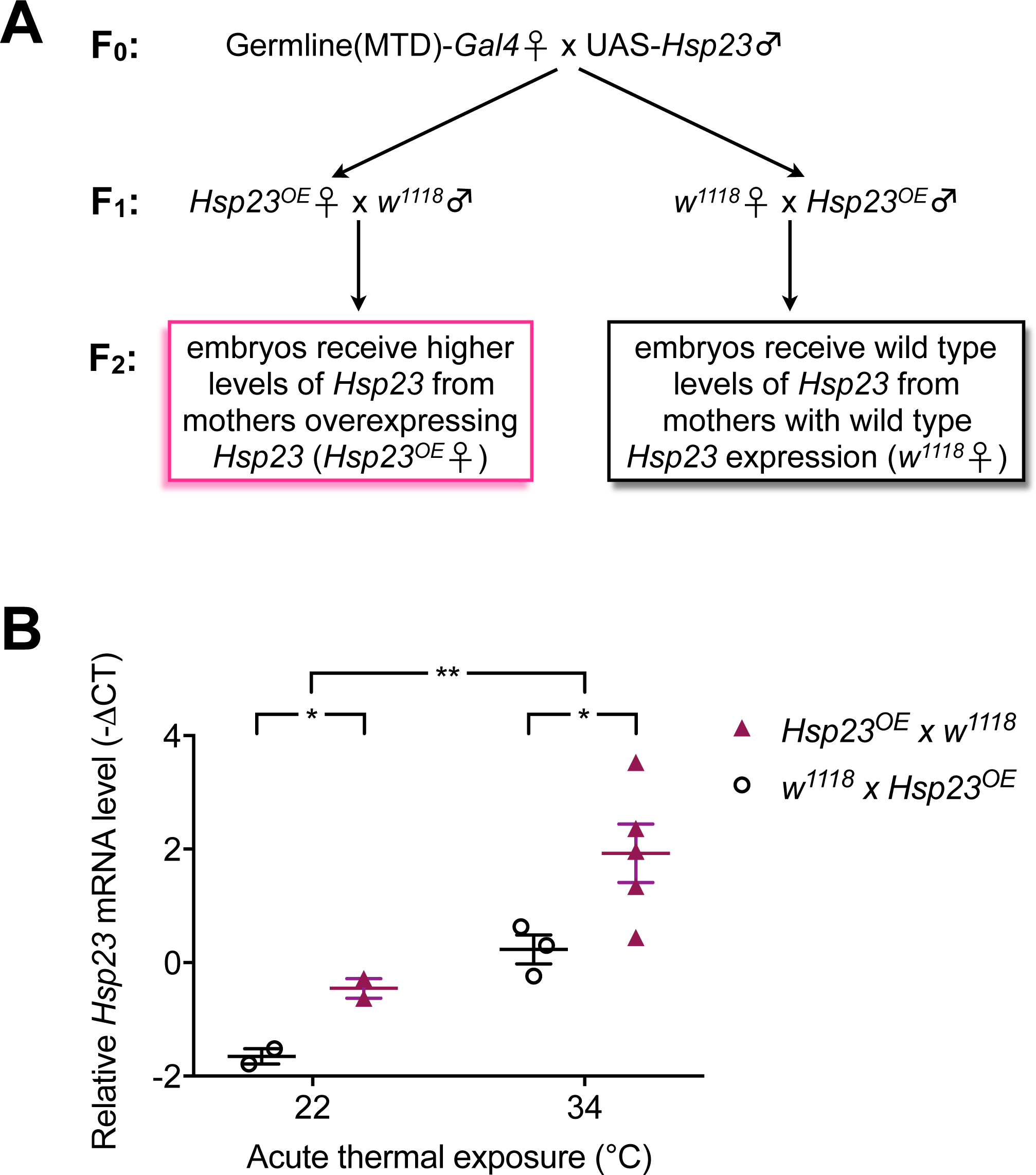
*Hsp23* overexpression in female ovaries increased *Hsp23* mRNA levels in offspring embryos. (A) Crossing scheme used to drive overexpression of *Hsp23* in the maternal germline, thus increasing maternal loading of *Hsp23* mRNAs into offspring embryos. The genotype of each sex is indicated by the gender symbols (♀ female x ♂ male). The MTD-*Gal4* construct drives overexpression of the UAS target gene (*Hsp23*) in the germline of F1 females (*Hsp23^OE^* ♀) but not males (*Hsp23^OE^* ♂). Reciprocal crosses between *Hsp23^OE^* and the control genetic background (*w^1118^*) produce F2 embryos that possess differential levels of maternally loaded *Hsp23*, as shown in part B. (B) Relative *Hsp23* mRNA levels in early embryos, normalized to the *Act5c* reference gene for each sample, expressed as -ΔCT, where ΔCT = CT*_Hsp23_*– CT*_Act5c_*. Each data point represents a pooled batch of 50 to 100 embryos (0-1 h-old) that were exposed to the indicated temperature for 45 min. Horizontal lines indicate means among separate embryo batches and error bars indicate standard error. \**P* < 0.05, \*\**P* < 0.01 (ANOVA temperature effect, *F*_1,8_ = 16.45, *P* = 0.0037, maternal genotype effect, *F*_1,8_ = 7.572, *P* = 0.025, temperature x maternal genotype interaction, *F*_1,8_ = 0.2216, *P* = 0.6504).

We focused our experiments on the maternal effects of overexpression and increased loading of *Hsp23* and did not perform a targeted knockdown of this gene for the following reasons. First, *Hsp23* is present in such low abundance in early embryos (Table 1; Fig. 1) that knocking down the expression of this gene is likely to have little effect. Additionally, there is evidence that the more abundant sHSPs, such as *Hsp26* and *Hsp27*, compensate for the absence of *Hsp23* under heat stress conditions (Bettencourt et al. 2008). Second, recent reviews of the literature suggest that, despite the preponderance of targeted gene knockdown experimental designs, many loss-of-function studies across a broad array of species have failed to produce measureable phenotypic outcomes (Gibney et al., 2013; Evans, 2015). This may be due to functional redundancy among genes or lack of assay sensitivity to characterize more subtle physiological effects (Bischof et al., 2013). Whatever the biological significance of these trends in loss-of-function studies, gain-of-function experimental designs are warranted and have led to the recent creation of comprehensive genetic resources for targeted gene over-expression (Bischof et al., 2013). Third, we predicted that overexpression and increased maternal loading of *Hsp23* into early embryos would more closely phenocopy the higher thermal tolerance of later stages of development that possess an enhanced ability, relative to early embryos, to induce the high levels of expression of heat shock genes, including *Hsp23* (Fig. 1).

**Table 1.** mRNA levels of sHSP genes in 5 day-old adults, 0-2 h-old embryos, and in response to heat-shock (4 day-old adults at 36°C for 1 h). Data are expressed as mean expression values (RPKM) and are ordered according to expression levels in heat-shocked adults. Data are from modENCODE (Brown et al. 2014).

| CG number | Gene Name | Adult Male<br>(5 d) | Adult Female<br>(5 d) | Embryo<br>(0-2 h) | Heat-shock<br>(Adult - 4 d) |
| --- | --- | --- | --- | --- | --- |
| CG4183 | *Hsp26* | 134 | 751 | 1276 | 3792 |
| CG4466 | *Hsp27* | 57 | 224 | 515 | 2689 |
| CG4463 | *Hsp23* | 43 | 13 | 25 | 2604 |
| CG4460 | *Hsp22* | 18 | 7 | 15 | 635 |
| CG4533 | *l(2)efl* | 209 | 63 | 2 | 257 |
| CG14207 | - | 115 | 55 | 50 | 219 |
| CG7409 | - | 421 | 84 | 0 | 76 |
| CG4190 | *Hsp67Bc* | 3 | 0 | 0 | 48 |
| CG4461 | - | 304 | 66 | 0 | 43 |
| CG4167 | *Hsp67Ba* | 1 | 0 | 0 | 16 |
| CG13133 | - | 8 | 1 | 0 | 1 |
| CG43851 | - | 4 | 0 | 0 | 1 |

### Quantification of *Hsp23* mRNA levels

We extracted total RNA from separate pooled batches of 20 - 100 embryos (0-1 h-old) that constituted our biological replicates for quantitative PCR (qPCR). Embryos were collected from grape juice agar plates after being exposed to 22°C or 34°C for 45 min (see Embryonic thermal tolerance section below), rinsed in 1x PBS, dechorionated in 50% bleach for 1 min, and rinsed again in diH_2_O. We note that embryos were less than 2 h-old, which is prior to the activation of zygotic transcription of the majority of genes (Ali-Murthy et al., 2013; Blythe and Wieschaus, 2015b). The embryos were then transferred to microcentrifuge tubes, frozen on liquid nitrogen, and stored at −80°C for up to one month prior to RNA extraction. We extracted total RNA with TRIzol (Molecular Research Center, Cincinnati, OH, USA) and Phase Lock Gel tubes (Quantabio, Beverly, MA, USA) that are designed to maintain stable separation of aqueous and organic phases. RNA quality was assessed on a NanoDrop spectrophotometer (NanoDrop products, Wilmington, DE, USA). We then removed any residual DNA with the TURBO DNA-FREE kit (Thermo Fisher Scientific, Waltham, MA, USA). We performed reverse transcriptase reactions with the SuperScript III First-Strand Synthesis kit using oligo dT primers (Thermo Fisher). qPCR was conducted with the Agilent Brilliant III Ultra-Fast SYBR Green Master Mix (Agilent Technologies, Santa Clara, CA, USA) on a Bio-Rad CFX Connect Real-Time PCR system (Bio-Rad, Hercules, CA, USA) using primer sets and reaction conditions that have been previously described (Bettencourt et al., 2008).

We calculated reaction efficiencies from standard curves for both the target gene (*Hsp23*) and the reference (*Act5c*), and we found the efficiencies to be identical for both genes (E = 1.87). We chose *Act5c* as the reference gene based on previous work that has shown it to exhibit stable expression across benign and heat shock conditions (Hoekstra and Montooth, 2013). We used the efficiency value to calculate average fold-differences among experimental groups as previously described (Pfaffl, 2001). We compared relative *Hsp23* mRNA levels among experimental groups using an ANOVA of -ΔCT values, where ΔCT = CT*_Hsp23_* – CT*_Act5c_*.

### Embryonic thermal tolerance

We measured acute thermal tolerance in early embryos (0 to 1 h post-fertilization) from crosses between genotypes designed to generate overexpression or normal expression (see above)(Fig. 2A). At this early stage, embryos coordinate early developmental processes via the molecular factors (i.e. RNA and protein) provided to them by their mothers (Tadros and Lipshitz, 2009; Blythe and Wieschaus, 2015a). Thus, embryonic phenotypic effects that we report on herein are largely the result of maternal effects mediated by changes in the expression of genes in female ovaries, the products of which are subsequently loaded into eggs. We chose this early stage because we sought to characterize the effects of (1) maternal mRNA contributions and (2) targeted gene overexpression in the absence of a fully developed zygotic heat-shock transcriptional response. This allowed us to better isolate and characterize the functional contribution of the transcription of a single gene (i.e. *Hsp23*) for whole-organism acute thermal tolerance, without the potentially confounding effects of large and concomitant changes in the expression of other heat shock genes.

We designed our temperature treatments to mimic sudden (acute) changes in temperature that frequently occur in nature where the temperature of necrotic fruit can increase rapidly on a hot day (Feder et al., 1997; Terblanche et al., 2011). 3 to 5 day-old adult flies of the appropriate genotypes were allowed to mate and lay eggs on grape juice agar plates for 1 h at 22°C. Egg plates were then wrapped in Parafilm and submerged in a water bath set to one of a range of temperatures between 22°C and 40°C (22°C, 24°C, 26°C, 28°C, 30°C, 32°C, 34°C, 36°C, 38°C, or 40°C) for 45 minutes. Due to the thermal mass of the egg plates, the embryos did not immediately experience the temperature of the water bath upon immersion, but rather were exposed to a thermal ramp that averaged +0.4°C min^-1^ among all temperatures. While this rate of change is extreme, it is within the range of maximum measured rates of change in the field (Terblanche et al., 2011). After thermal exposure, a section of the agar containing 20 eggs was cut and transferred to a food vial where eggs were allowed to recover and develop at 22°C. Hatching, pupation, and eclosion success were scored as the proportion of these 20 eggs that survived to each stage. Hatching success was scored at 48 h, pupation success at 5 to 10 days, and eclosion at 10 to 15 days post-fertilization. We also scored development time as the length of time (days) to successful pupation and eclosion. Temperature treatments and phenotypic measurements were conducted on four to six vials in each of three to four separate generations for each cross type (i.e., N = 4 to 6 vials x 10 temperatures x 3 to 4 generations per genotype) for a total number of 12 to 24 biological replicates per genotype per temperature.

We calculated the lethal temperature at which 50% of the embryos failed to hatch, pupate, or eclose (LT_50_) via a least-squares logistic regression model. We allowed the y-intercept to vary between 0 and 1 and extrapolated the LT_50_ from the inflection point of the logistic curve fit. This approach allowed us to infer thermal tolerance independently from other confounding factors that reduce hatching success, such as the presence of unfertilized eggs.

### Pupation height

We scored the average pupation height as a measurement of larval performance (Mueller and Sweet, 1986; Hoekstra et al., 2013). Pupation height was scored subsequent to early embryonic temperature treatments (see above) at 8 to 10 days post-fertilization. Each food vial was divided into four quadrants; quadrant 1 spanned the distance from the bottom of the vial to 3.5 cm in height and quadrants 2 - 4 each comprised a section of 2 cm up the height of the vial. Each pupa was scored a number between 1 and 4, corresponding to the quadrant in which it pupated. All pupae on the food were scored as 1 (quadrant 1). Average pupation height was then calculated separately for each vial.

### Statistics

LT_50_ values were compared by assessing the fit of the logistic regression models to each genotype separately vs. all genotypes combined via the extra sum-of-squares F-test and the Akaike’s Information Criteria (AICc). The effects of temperature, treatment, and maternal genotype on development time were analyzed via ANOVA followed by Sidak’s multiple comparisons test to assess pairwise differences. Pupation height was analyzed in the same manner via ANOVA and Sidak’s test. All analyses were conducted in GraphPad Prism version 7 for Mac (GraphPad Software, La Jolla, CA).

## RESULTS

Among all 18,029 transcripts included in the modENCODE dataset, mRNA levels in early embryos and heat-shocked adults were positively correlated (Fig. 1; Least-squares regression, *R^2^* = 0.17, *y* = 10^(1.002*log(*x*) - 0.1344)). Among the 12 sHSP genes, mRNA levels in embryos and heat-shocked adults were also positively correlated (Fig. 1; Robust regression, *R^2^* = 0.97, *y* = 10^(0.5436*log(*x*) + 1.905)), even though mRNA levels were higher in heat-shocked adults compared to embryos (Table 1; Fig. 1). However, *Hsp23* was a significant outlier in this relationship (Fig. 1; ROUT outlier analysis, Q = 1%). Of sHSP genes, *Hsp23* has the biggest difference in expression level between early embryos and heat-shocked adults and is present at low levels in non-heat-shocked adults, with a heat-shock induction response of >100-fold (Table 1).

We sought to test the contribution of maternal *Hsp23* mRNAs to embryonic thermal tolerance by increasing *Hsp23* abundance in early embryos through overexpression in the maternal germline. We focused our functional genetic analyses on *Hsp23* because this gene (1) was the sole significant outlier among the sHSP genes in the relationship between early embryonic vs. heat-shocked adult gene expression (Fig. 1) and (2) showed the greatest induction in response to heat-shock in adults (Table 1). These observations suggest that *Hsp23* plays a unique role among the sHSP genes in the heat-shock response, and thus may be a key factor in conferring acute thermal tolerance.

Maternal genotype (*Hsp23^OE^* versus *w^1118^* control) and embryonic heat stress (45 min at 34°C) both had significant effects on *Hsp23* mRNA levels in early embryos (Table 2 and Fig. 2B; ANOVA temperature effect, *F*_1,8_ = 16.45, *P* = 0.0037, maternal genotype effect, *F*_1,8_ = 7.572, *P* = 0.025), and these effects were additive (ANOVA temperature x maternal genotype interaction, *F*_1,8_ = 0.2216, *P* = 0.6504). *Hsp23*- overexpressing females (*Hsp23^OE^*) laid eggs with 2.12-fold higher baseline levels of *Hsp23* mRNAs at 22°C and a 2.89-fold higher levels of *Hsp23* mRNAs following heat shock at 34°C, relative to embryos that were offspring of mothers of the control genetic background (*w^1118^*) (Fig. 2**B**). In addition, heat shock led to significant increases in the levels of *Hsp23* mRNAs regardless of maternal genotype, increasing by 4.44-fold and 3.26-fold (34°C relative to 22°C) in offspring embryos of (female x male) *Hsp23^OE^* x *w^1118^* and *w^1118^* x *Hsp23^OE^*, respectively (Fig. 2B).

**Table 2.** Analysis of variance of relative *Hsp23* mRNA levels (ΔCT = CT*_Hsp23_* - CT*_Act5c_*) among maternal genotypes (*Hsp23^OE^* vs. *w^1118^*^)^ and embryonic heat stress temperatures (22°C vs. 34°C).

| Source of variation | SS | DF | MS | F (1,8) | % of total variation | P-value |
| --- | --- | --- | --- | --- | --- | --- |
| Temperature | 11.88 | 1 | 11.88 | 16.45 | 43.54 | 0.0037 |
| Maternal genotype | 5.467 | 1 | 5.467 | 7.572 | 20.04 | 0.025 |
| Temp. x Mat. genotype | 0.16 | 1 | 0.16 | 0.2216 | 0.59 | 0.65 |
| Residual | 5.776 | 8 | 0.722 |  |  |  |

Maternal overexpression of *Hsp23* significantly increased embryonic thermal tolerance by raising the lethal temperature (LT_50_) by approx. 1°C (Fig. 3; Extra sum of squares F-test, *F*_1,262_ = 5.371, *P =* 0.02). Embryos that successfully hatched also survived to pupation and adulthood, as 95-100% of larvae and pupae survived to pupation and eclosion, respectively, regardless of maternal genotype. Furthermore, there were no significant differences between the LT_50_ of hatching, pupation, and eclosion successes for a given genotype (Fig. 3B; Extra sum of squares F-test, *P* > 0.05), suggesting that effects of early, acute thermal stress on survival were largely localized to embryogenesis.

**Figure 3.**
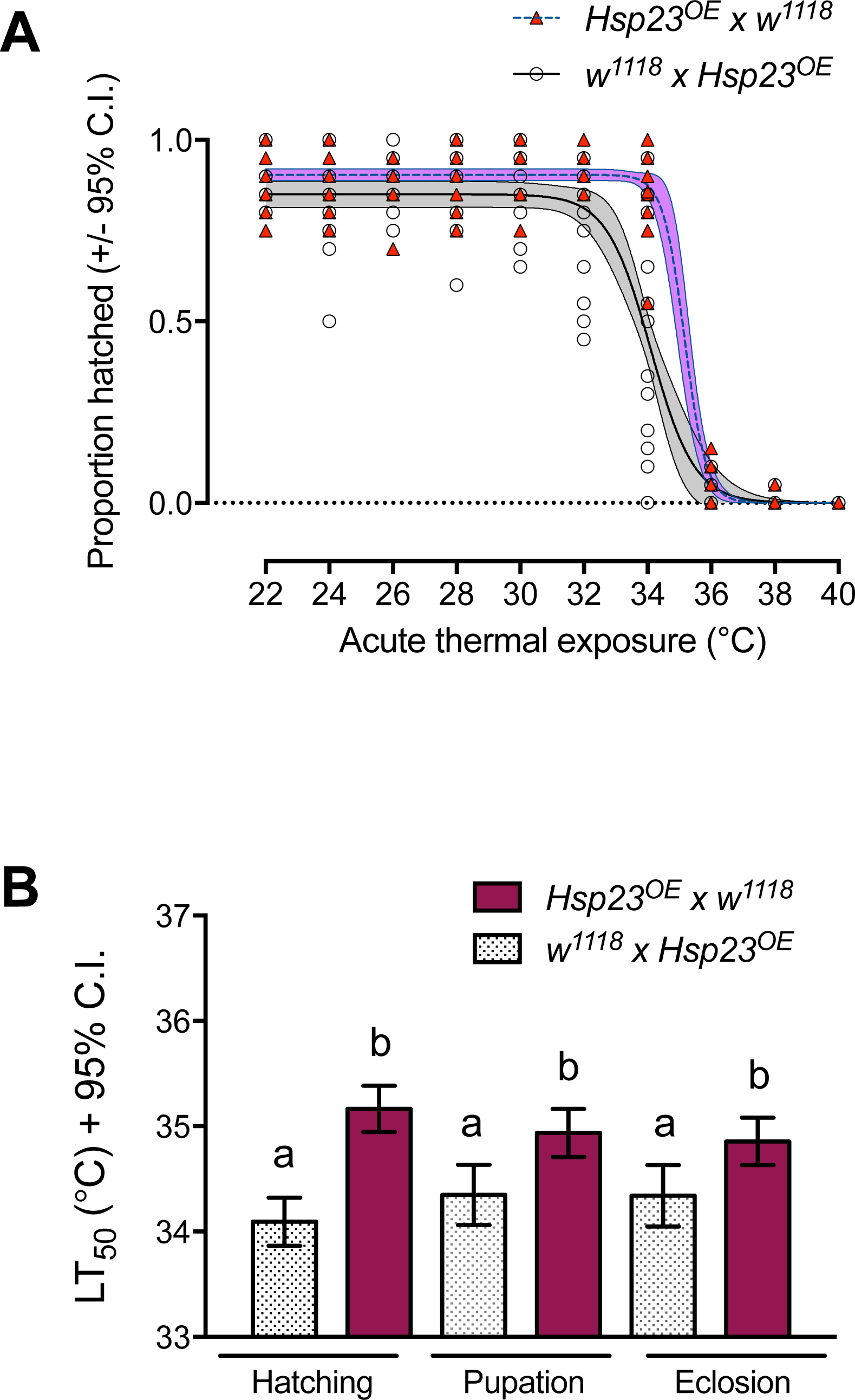
Higher maternal loading of *Hsp23* mRNAs increased thermal tolerance of offspring embryos. (A) Each data point represents the proportion of eggs (N = 20 eggs, 0-1 h-old) that hatched following 45 minutes exposure at the indicated temperature. Lines indicate least-squares fit of the logistic equation and shaded regions indicate 95% confidence bands. (B) Mean LT_50_ values +/- 95% confidence intervals for hatching, pupation, and eclosion success. LT_50_ values were extrapolated from the inflection points of the logistic survival curve fits (hatching success is shown in part A), and confidence intervals represent the goodness of fit of the logistic regressions. Different letters indicate statistical significance (*P* < 0.05, Extra sum of squares F-test). The genotypes of the parents are indicated in the legends of both figure panels (female x male).

In addition to the positive and protective effect of maternal *Hsp23* overexpression for whole-embryo survival of thermal stress, maternal loading of this gene in early embryos had significant effects on larval performance, as indexed by pupation height. Exposure of 0-1 h-old embryos to the brief (45 min.) episode of thermal exposure resulted in larvae with significantly reduced pupation height at the highest temperatures (Fig. 4A), which explained 22% of the variation in pupation height (Table 3; ANOVA temperature effect, *F*_8,189_ = 7.744, *P* < 0.0001). Maternal *Hsp23* overexpression had no significant effect on pupation height overall (Fig. 4A; Table 3; ANOVA maternal genotype effect, *F*_1,189_ = 3.674, *P* = 0.0568) but conferred protection against the negative effects of heat stress on pupation height, particularly at 34°C (Fig. 4A; Table 3; ANOVA temperature x maternal genotype interaction, *F*_8,189_ = 2.822, *P* = 0.0056, Sidak’s test on pairwise difference at 34°C, *P* < 0.001).

**Table 3.** Percent of total variation explained by the main effects of embryonic heat stress temperature, maternal genotype, and their interaction on pupation height and days to pupation (Two-way ANOVA). *P*-values are indicated in parentheses and significant effects are indicated in italics.

| Trait | Temperature | Maternal genotype | Temp. x Mat. gen. |
| --- | --- | --- | --- |
| Pupation height | *21.63% (< 0.0001)* | 1.28% (0.0568) | *7.88% (0.0056)* |
| Days to pupation | *28.97% (< 0.0001)* | 1.10% (0.0889) | 1.99% (0.73) |

Embryonic heat stress also caused significant increases in development time, as indexed by the length of time (days) to pupation (Fig. 4B), in offspring of both maternal genotypes, with a trend of maternal overexpression of *Hsp23* attenuating this developmental delay (Table 3; ANOVA temperature effect, *F*_8,169_ = 9.605, *P* < 0.0001, maternal genotype effect, *F*_1,169_ = 2.928, *P* = 0.0889, temperature x maternal genotype interaction, *F*_8,169_ = 0.6591, *P* =0.73).

**Figure 4.**
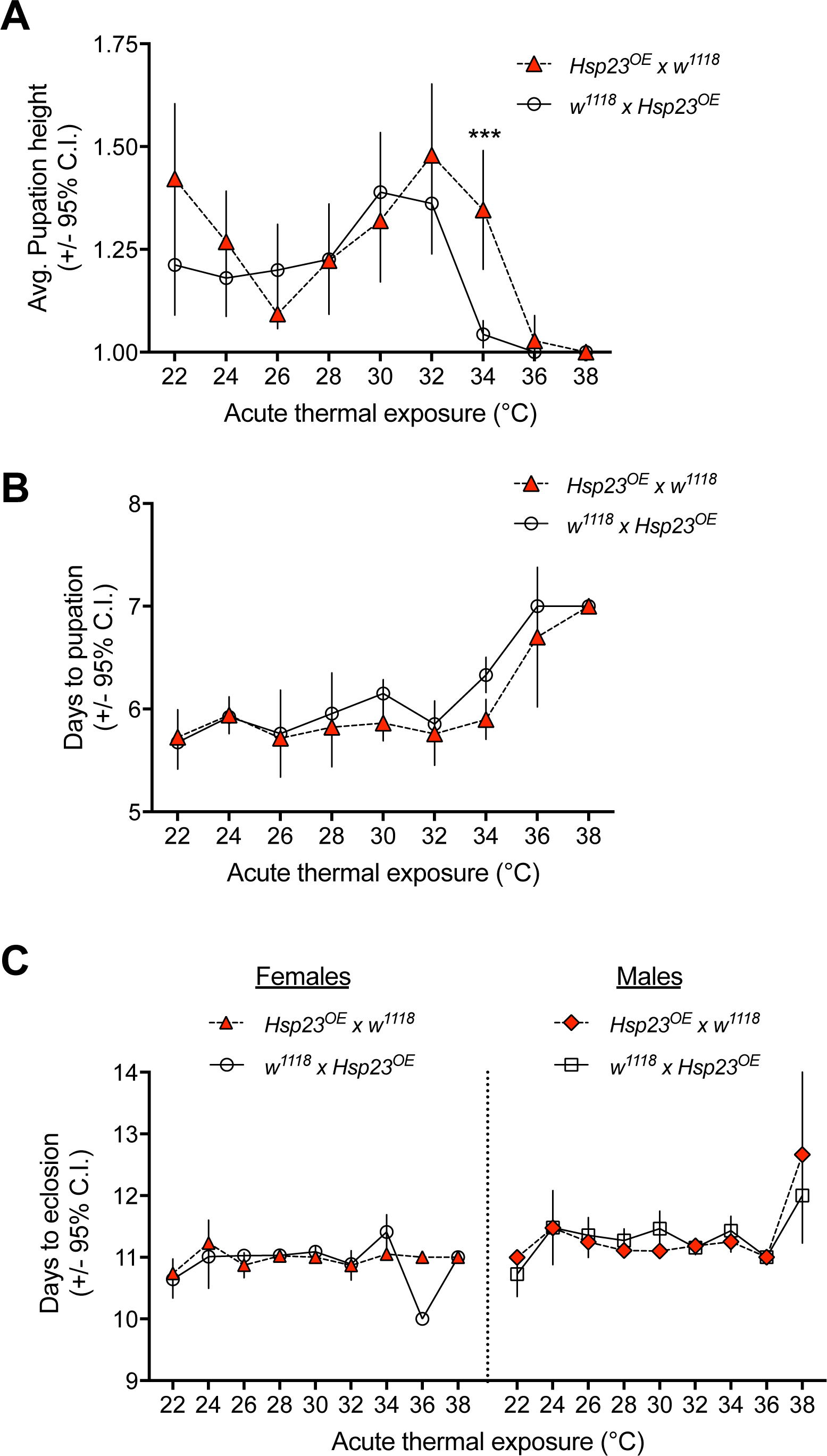
Heat stress in early embryos caused larval defects that were ameliorated by maternal loading of *Hsp23*. (A) Mean pupation height among vials (N = 18 vials), +/- 95% confidence intervals, scored at 5 to 10-days-post-fertilization following acute (45 min) early embryonic (0-1 h-post-fertilization) exposure at the indicated temperature. Higher heat stress temperatures caused significant decreases in pupation height (ANOVA temperature effect, *F*_8,189_ = 7.744, *P* < 0.0001), but higher maternal loading of *Hsp23* removed this effect at 34°C (Sidak’s test on pairwise difference at 34°C, \*\*\**P* < 0.001). (B) Mean time to pupation (days +/- 95% confidence intervals) following early embryonic temperature exposure, as described above in A. Higher embryonic temperature exposure caused increased time to pupation (Table 3; ANOVA temperature effect, *F*_8,169_ = 9.605, *P* < 0.0001), and there was a trend of maternal loading of *Hsp23* attenuating this effect (maternal genotype effect, *F*_1,169_ = 2.928, *P* = 0.0889, temperature x maternal genotype interaction, *F*_8,169_ = 0.6591, *P* =0.73). (C) Mean time to eclosion (days +/- 95% confidence intervals) following early embryonic temperature exposure, as described above in A. Males took longer to eclose than females at all temperatures (ANOVA, sex effect, *F*_1,347_ = 30.263, *P* < 0.00001), but the effects of heat stress on development time were similar between the sexes (ANOVA, temperature x sex interaction, *F*_8,347_ = 0.149, *P* = 0.70). Data represent values from females (left panel) and males (right panel). The genotypes of the parents are indicated in the legends of each panel (female x male).

Embryonic heat stress also significantly affected the developmental time to adult eclosion (Fig. 4C; Table 4); however, the effect sizes were much smaller than the heat-stress-induced delay to pupation, and the pattern was largely driven by a shorter time to eclosion at 22°C (Fig. 4C; ANOVA temperature effect, *F*_8,347_ = 11.511, *P* < 0.001). Maternal *Hsp23* overexpression had no significant effect on time to eclosion (Fig. 4C; Table 4; ANOVA maternal genotype effect, *F*_1,347_ = 0.814, *P* = 0.3676), regardless of temperature (ANOVA temperature x maternal genotype interaction, *F*_8,347_ = 2.020, *P* = 0.1561), sex (ANOVA sex x maternal genotype interaction, *F*_1,347_ = 0.098, *P* = 0.7544), or the interaction among all of these effects (ANOVA temperature x sex x maternal genotype interaction, *F*_1,347_ = 0.1091, *P* = 0.7414). There was a significant difference between females and males in time to eclosion across all temperatures, with females eclosing sooner than males, and sex accounted for the greatest variation in time to eclosion (Table 4; Fig. 4C; ANOVA sex effect, *F*_1,347_ = 30.263, *P* < 0.00001). And, while males on average suffered greater developmental delays to eclosion following acute exposure to 38°C (Fig. 4C), this effect was not significant (ANOVA temperature x sex interaction, *F*_8,347_ = 0.149, *P* = 0.70).

**Table 4.** Percent of total variation explained by the main effects of embryonic heat stress temperature, maternal genotype, sex, and their interaction on days to adult eclosion (3-way ANOVA). *P*-values are indicated in parentheses and significant effects are indicated in italics.

| Source of variation | % Variation explained (*P*-value) |
| --- | --- |
| Temperature | *2.94% (< 0.001)* |
| Maternal genotype | 0.21% (0.3676) |
| Sex | *7.72% (< 0.00001)* |
| Temp. x Mat. genotype | 0.52% (0.1561) |
| Temp. x Sex | 0.038% (0.70) |
| Mat. genotype x Sex | 0.025% (0.7544) |
| Temp x Mat. genotype x Sex | 0.028% (0.7414) |

There was a slight discrepancy between the thermally induced delays in development to pupation vs. eclosion. Specifically, embryonic thermal stress at 34°C and 36°C caused delays in time to pupation but not eclosion (Figs. 4B and 4C). In effect, this means that the pupae that suffered developmental delays to pupation somehow recovered from this delay to be able to eclose on the same schedule as pupae that were exposed to lower embryonic temperatures. This may have occurred due to the entrainment of eclosion behavior by circadian rhythms (Kyriacou et al., 1990; Paranjpe et al., 2005), in which case delayed pupae could catch up to the eclosion schedule of other pupae, as long as the pupation was delayed by less than 24 hours. Alternatively, this pattern may be an artifact of the low sample sizes and high variance in development times that accompanied the more extreme thermal exposures, as far fewer individuals successfully hatched after exposure to the highest temperatures (Fig. 3A). But despite this incongruity between pupation and eclosion times at the highest temperatures of embryonic heat stress, overall our data indicate that overexpression of *Hsp23* in the maternal germline not only increased embryonic hatching success after exposure to heat stress, but also had persisting effects on offspring performance throughout larval development.

## DISCUSSION

Despite over five decades of research on the heat shock response, there have been relatively few studies to connect genotype to phenotype in the context of heat shock protein expression and organismal performance (Somero et al., 2017). Here we demonstrate the direct effects of maternal loading of *Hsp23* mRNAs for offspring survival and performance following acute heat stress in a common genetic background. We found that increases in the levels of *Hsp23* in early *D. melanogaster* embryos confer significant protection from heat stress during a thermally sensitive life stage.

### *Hsp23* maternal loading and the embryonic heat shock response

*Hsp23* overexpression in female ovaries resulted in embryos with increased abundance of *Hsp23* mRNA. Thus, the phenotypic effects of maternal genotype on whole-embryo thermal tolerance that we observed were likely the consequence of increased maternal loading that elevated basal levels of *Hsp23* in early embryos. We also observed that embryos induced the expression of *Hsp23* in response to heat shock regardless of maternal genotype, with no significant interaction between maternal genotype and temperature. Previous work has shown that the transgenic manipulation of heat shock protein 70 (*Hsp70*) gene induction in *D. melanogaster* (Welte et al., 1993) causes massive increases in the heat-induced transcription of *Hsp70* by more than 500- fold (Hoekstra and Montooth, 2013). This increase in heat-inducible *Hsp70* mRNAs translated into approximately 2.5-fold higher levels of Hsp70 protein over the time course of two hours in response to heat stress in wandering third-instar larvae, which allowed larvae to survive significantly longer at 39°C (Feder et al., 1996). In comparison, the higher levels of *Hsp23* induced by Gal4/UAS overexpression in female ovaries that we report herein were subtle. These overexpression levels were similar to previous reports of overexpression of other genes in fly ovaries that were driven by similar transgenic constructs (Dominguez et al., 2016). But regardless of the absolute degree of overexpression, the higher maternal loading of *Hsp23* increased embryonic LT_50_ by approx. 1°C, and this increase in LT_50_ signified a substantial increase in thermal tolerance. In particular, following 45 minutes of heat stress at 34°C, 87.5% of the embryos with higher *Hsp23* levels successfully hatched, whereas only 46.7% of embryos with normal levels of *Hsp23* survived this heat treatment.

Beyond the aforementioned work in *D. melanogaster* (Welte et al., 1993; Feder et al., 1996) and the present study, there have been few studies to directly test the effect of heat shock protein expression on whole-organism thermal tolerance, with one study showing that targeted gene knockdown of *Hsp22* and *Hsp23* in adult *D. melanogaster* decreases cold tolerance (Colinet et al., 2010). A much larger body of work has used interspecies and inter-population comparisons to infer the evolutionary history of the heat shock response (Hofmann and Somero, 1996; Tomanek and Somero, 2000; Dong et al., 2008; Lockwood et al., 2010; Schoville et al., 2012; Dowd and Somero, 2013; Nguyen et al., 2016) Based on these studies, it is well-established that HSP expression is an adaptive physiological mechanism for coping with acute thermal stress. Accordingly, population-level comparisons have discovered clinal variation in *Hsp23* alleles across environmental thermal gradients in *Drosophila buzzatii* in Australia (Frydenberg et al., 2010), as well as clines in allele frequencies of *Hsp23* and *Hsp26* among *D. melanogaster* populations in Australia (Frydenberg et al., 2003). In addition, laboratory thermal selection studies have found evolved changes in HSP expression to accompany adaptive shifts in upper thermal limits (Rudolph et al., 2010; Kelly et al., 2017). It is important to note, however, that increased levels of HSP expression do not always accompany thermal adaptation to higher temperatures (Zatsepina et al., 2001). In fact, experimental evolution to a higher constant temperature in *D. melanogaster* led to the evolution of lower Hsp70 expression and concomitant decreases in acute thermal tolerance (Bettencourt et al., 1999). Thus, higher basal and inducible HSP expression may be adaptive primarily in environments that are characterized by sudden and dramatic heat stress events, rather than constant hot environments (Dong et al., 2008; Dilly et al., 2012).

Even though our observed genotypic effects on embryo thermal tolerance were the result of differential maternal loading of *Hsp23*, it is interesting to note that we observed zygotic induction of this gene in offspring of both maternal genotypes. At this early stage of development (0-1 h-old), zygotic genomes are predicted to be transcriptionally inactive because embryos have not undergone the maternal-to-zygotic transition (MZT) that occurs in the mid-blastula stage (approx. 2.5 h-old) in *D. melanogaster* (Tadros and Lipshitz, 2009; Blythe and Wieschaus, 2015a). However, prior to the canonical MZT, zygotic gene expression appears to be responsive to thermal variability. A previous analysis of protein expression in early *D. melanogaster* embryos using 2-dimensional gel electrophoresis found that heat shock proteins were heat-inducible at 1 to 2-h post-fertilization (Graziosi et al., 1980). Moreover, recent work in *D. melanogaster* has highlighted the developmental role of early zygotic gene transcription that precedes the MZT (Ali-Murthy et al., 2013), but the full extent to which the early zygotic genome responds to thermal variability warrants new investigation. In the present study, *Hsp23* expression was induced in early embryos to a much lesser extent (approx. 4-fold) than what has been previously observed in later stages of development. In fact, Leemans et al. (Leemans et al., 2000) found *Hsp23* to be heat-induced by more than 10-fold in late-stage embryos (18 h-old) and Brown et al. (Brown et al., 2014) reported this gene to be heat-induced by approx. 100-fold in adults (Table 1). Therefore, while embryos at the earliest stages of development appear to exhibit a heat shock response, it is at a much-reduced level compared to later stages. This explains why early embryonic stages are more thermally sensitive than later stages (Walter et al., 1990; Welte et al., 1993) and further emphasizes the potentially critical role of maternally-loaded mRNAs and proteins as thermal protectants.

The specific mechanism by which the Hsp23 protein confers thermal tolerance remains elusive. This protein exhibits general chaperoning activity by preventing heat denaturation of proteins in vitro (Heikkila et al., 2006), but it has also been shown to be involved in ventral furrow morphogenesis in early fly embryos under benign thermal conditions (Gong et al., 2004). This developmental role may be mediated through the interaction of Hsp23 with elements of the cytoskeleton, such as microtubules (Hughes et al., 2008) and actin microfilaments (Goldstein and Gunawardena, 2000). The cytoskeletal association of Hsp23 is further supported by the observation that this protein was the only small heat shock protein whose overexpression was observed to prevent actin-dependent contractile dysfunction in cardiomyocytes of *D. melanogaster* larvae (Zhang et al., 2011). Indeed, among the sHSPs that are highly induced in response to heat stress (Table 1), Hsp23 is the only one that is both localized to the cytoplasm (Morrow and Tanguay, 2015) and contains an actin-binding domain (sequence data not shown). Whether or not the interaction of Hsp23 with the cytoskeleton provides protection in the context of thermal stress has not been reported, but this is a worthwhile topic of future study.

### Effects of embryonic heat stress on post-embryonic larval development

We found that the *Hsp23*-mediated maternal effect extended beyond embryonic thermal tolerance (i.e., hatching success) and attenuated heat-induced defects in larval performance (i.e., pupation height) and prevented heat-induced developmental delays (i.e., days to pupation). It is surprising that a brief thermal exposure experienced during the first 2 hours of life has negative consequences that last for days to weeks, throughout larval development. Drops in pupation height and increases in development time have been previously associated with lower energetic performance and fitness (Mueller and Sweet, 1986; Montooth et al., 2010; Hoekstra et al., 2013; Meiklejohn et al., 2013), and may have important ecological consequences in natural populations.

The persistent effects of maternal transcript loading on larval development post-heat stress might have important consequences for the evolution of maternal effects. A recent study reported significant maternal effects that determined both acute (i.e., 1 h at 27°C) and chronic (i.e., constant exposure to 24°C) thermal tolerance in offspring embryos among wild populations of *Ciona intestinalis* (Sato et al., 2015). This suggests that there is natural genetic variation for maternal effects of thermal traits in some species. However, evolutionary theory predicts that selection is less effective on alleles that confer maternal effects, compared to genes expressed in both sexes, due to a reduced effective population size (Demuth and Wade, 2007; Van Dyken and Wade, 2010). Consequently, maternal effect genes can harbor higher levels of standing genetic variation, presumably because deleterious mutations are not removed as frequently from the population and these genes cannot evolve as readily via positive selection (Barker et al., 2005). Nevertheless, if maternal effects not only determine hatching success but also influence larval performance and development, then the fitness consequences associated with thermal stress may lead to a greater strength of selection on maternal effect genes (i.e., greater difference in fitness among maternal effect genotypes) than would otherwise be predicted from the maternal effects of offspring hatching success alone. Because responses to natural selection depend on both the strength and the efficacy of selection, the broad developmental effects of maternal transcript loading may favor the adaptive evolution of maternal effect thermal traits, depending on the thermal environment (Chevin et al., 2010; Chevin and Hoffmann, 2016) and the underlying genetic architecture (Wolf and Wade, 2016). But to our knowledge, there have been no examples of this phenomenon reported in the literature.

### Conclusions

Overall, our data suggest that maternal effects can have profound impacts on offspring survival and performance in the context of environmental change. The observation that differential maternal loading of mRNAs of a single gene can have lasting consequences throughout larval development, by modifying pupation height and development time, demonstrates that protective maternal effects extend well beyond the maternal-to-zygotic transcriptional transition. The role of maternal effects and the environmental stress physiology of early life stages has largely been ignored in the field of ecological physiology, but see (Sato et al., 2015). Future work is warranted in this realm, because these factors are likely be critical determinants of species responses to environmental variability (Angilletta et al., 2013; Anderson and Podrabsky, 2014; Buckley et al., 2015; Wagner and Podrabsky, 2015; Svetec et al., 2016), particularly if early life stages are most sensitive to environmental stress (Walter et al., 1990; Welte et al., 1993).

## Acknowledgements

We thank Rosemary Scavotto and Sarah Howe for assistance with fly husbandry and Tarun Gupta for valuable discussions that aided in the preparation of the manuscript.

## Author contributions

B.L.L. and K.L.M. conceived and planned the study. B.L.L. and C.R.J. conducted the experiments. B.L.L. analyzed the data. B.L.L., C.R.J., and K.L.M. wrote the manuscript.

## Data availability

modENCODE transcript data are publicly available online at flybase.org (Gramates et al., 2017). Phenotypic data reported herein are available on the Dryad Repository, http://.

